# Cytosolic ROS production by NADPH oxidase 2 regulates muscle glucose uptake during exercise

**DOI:** 10.1101/522805

**Authors:** Carlos Henríquez-Olguin, Jonas R. Knudsen, Steffen H. Raun, Zhencheng Li, Emilie Dalbram, Jonas T. Treebak, Lykke Sylow, Rikard Holmdahl, Erik A. Richter, Enrique Jaimovich, Thomas E. Jensen

## Abstract

Reactive oxygen species (ROS) act as intracellular compartmentalized second messengers mediating metabolic stress-adaptation. In skeletal muscle fibers, ROS have been suggested to stimulate glucose transporter 4 (GLUT4)-dependent glucose transport during artificially evoked contraction *ex vivo* but whether myocellular ROS production is stimulated by *in vivo* exercise to control metabolism is unclear. Here, we combined exercise in humans and mice with fluorescent dyes, genetically-encoded biosensors, and NADPH oxidase 2 (NOX2) loss-of-function models to demonstrate that NOX2 is the main source of cytosolic ROS during moderate-intensity exercise in skeletal muscle. Furthermore, two NOX2 loss-of-function mouse models lacking either p47phox or Rac1 presented striking phenotypic similarities, including greatly reduced exercise-stimulated glucose uptake and GLUT4 translocation. These findings indicate that NOX2 is a major myocellular ROS source regulating glucose transport capacity during moderate-intensity exercise.

## INTRODUCTION

A single bout of exercise prompts a rapid adaptive increase in energy substrate metabolism in contracting skeletal muscle to meet the increased demand for ATP production. Understanding the molecular signal transduction pathways that orchestrate these metabolic responses in muscle has broad implications, from optimizing athletic performance, to understanding fundamental stress-adaptive responses at the cellular level, to prevention and treatment of aging- and lifestyle-related diseases in muscle and other tissues^1^.

The production of oxygen-derived free radicals and derivatives thereof, collectively referred to as reactive oxygen species (ROS), increases in skeletal muscle during exercise and has long been considered to mediate adaptive responses to both acute bouts of exercise and chronic exercise training^2^. Although historically viewed as a by-product of oxidative metabolism in mitochondria, it has been suggested that ROS may be produced enzymatically by extra-mitochondrial sources in contracting muscle, including NADPH oxidase (NOX)^3^ and xanthine oxidase^4^. However, the primary ROS source in the context of physiological *in vivo* exercise remains uncertain due to the difficulty in measuring and quantifying exercise-stimulated ROS production^5^.

Research in the past 60 years has established that glucose transport *in vivo* is controlled by a coordinated increase in glucose delivery, glucose transport into muscle fibers and intracellular metabolism^6, 7^ Among these events, the insulin-independent translocation of glucose transporter protein 4 (GLUT4) from intracellular storage depots to the cell surface to facilitate glucose entry appears to be a key molecular event^8, 9^. Studies in isolated *ex vivo* incubated skeletal muscles from rodents have shown that glucose transport in response to electrically stimulated contraction and mechanical stress is lowered by antioxidants^10, 11^, suggesting that ROS are required for increased glucose transport^12^. Interestingly, the small GTPase Rac1 is activated by muscle contraction and passive stretch and is necessary for both stimuli to increase glucose uptake in isolated mouse muscles^13, 14^. Rac1 is best known to bind and orchestrate regulators of actin remodeling^15^, but the recruitment of GTP-loaded active Rac GTPase isoforms in conjunction with a complex of other NOX2 regulatory proteins (p67phox, p47phox and p40phox) is also essential to stimulate superoxide (O_2_^−^)-production by the membrane-bound NOX2 complex^16^. However, whether NOX2-induced ROS production regulates muscle glucose uptake *in vivo* and if the reduced exercise-stimulated glucose uptake observed in Rac1 deficient muscles^17^ is due to reduced ROS production, is currently unknown.

In the present study, we took advantage of recent methodological developments in the redox signaling field, including a method for preservation of *in vivo* ROS modifications and genetically encoded ROS biosensors^18^. This allowed us to measure both general and localized NOX2-specific ROS production in wild-type (WT) mice and mice lacking NOX2 activity due to the absence of either the Rac1 or p47phox regulatory subunits. Using these approaches, we show that NOX2 is activated by moderate-intensity exercise and is the predominant source of ROS production under these conditions. Moreover, a large reduction in exercise-stimulated glucose uptake and GLUT4 translocation were shared features between the two NOX2 loss-of-function models. Collectively, these results imply that NOX2 is a major source of ROS production during moderate-intensity exercise and that Rac1 is required for GLUT4 translocation and glucose uptake due to its essential role in NOX2 activation.

## RESULTS

### Moderate-intensity exercise causes a pro-oxidative shift in human muscle

Plasma redox markers are increased during both moderate- and high-intensity exercise in humans^19^. However, whether muscle ROS production increases during moderate-intensity exercise in humans is unknown. To explore this, we estimated total oxidant production in skeletal muscle from healthy young men before and after a single 30 min bout of bicycle ergometer exercise at a moderate 65% peak power output exercise intensity using the redox-sensitive dye dichlorodihydrofluorescein diacetate (DCFH). We found that exercise stimulated an 86% increase in DCFH oxidation in human skeletal muscle (Fig. 1A), which was accompanied by an expected increase in the phosphorylation state of known exercise-responsive proteins (Fig. 1B). No changes in total protein expression were observed (Supplementary Fig. 1A) This shows that moderate-intensity exercise is pro-oxidative in human skeletal muscle.

**Figure 1.**
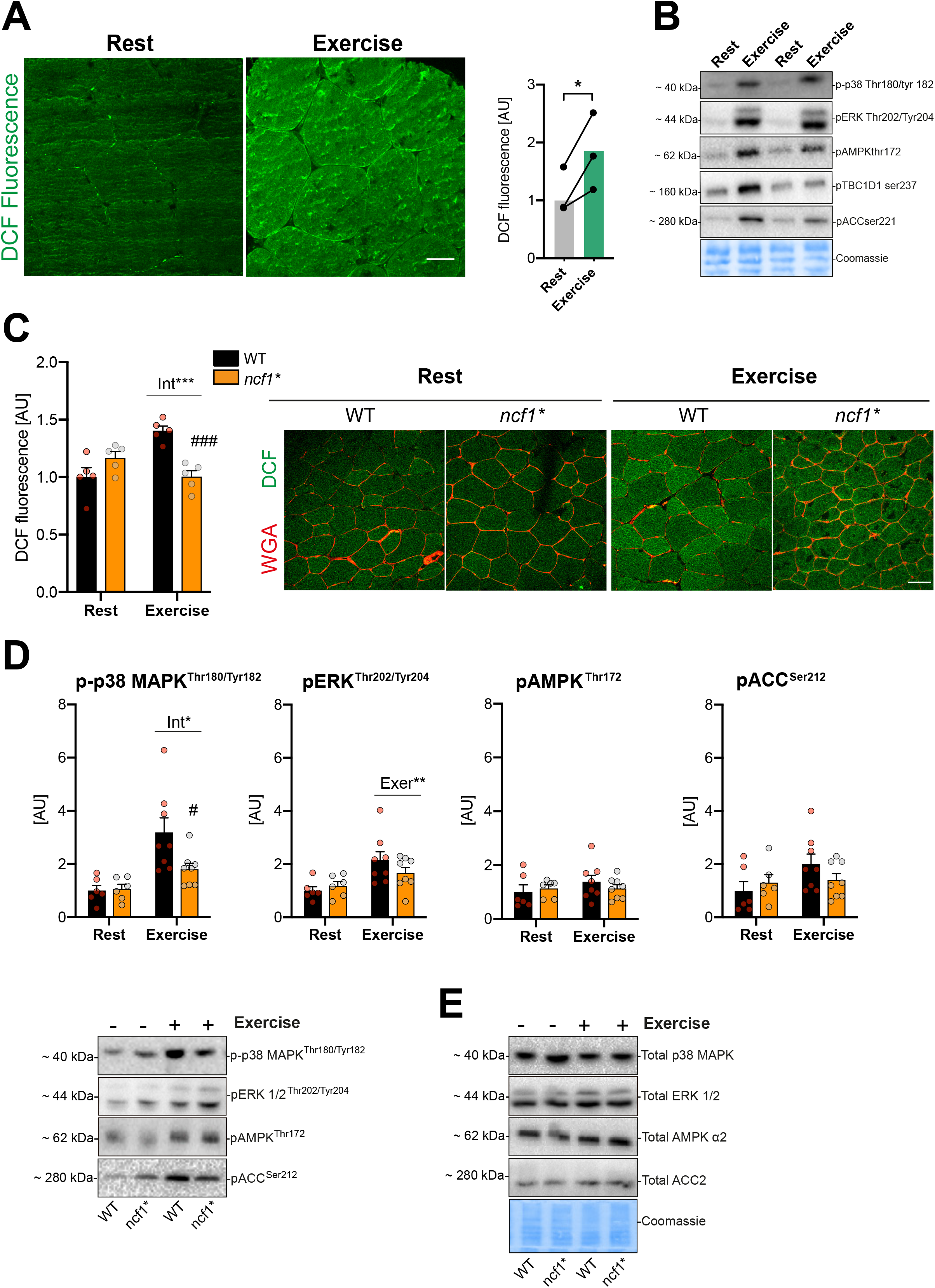
Moderate-intensity exercise causes a pro-oxidative shift in human and murine muscle. **a** 2’,7’-dichlorodihydrofluorescein diacetate (DCFH) oxidation (DCF) and **b** exercise signaling in human vastus lateralis before and after a bout of moderate-intensity cycling (30 min, 65% maximal power output) in young-healthy volunteers (n=3). **c** Representative images and quantification of the exercise-stimulated (20 min, 65% maximal running speed) DCFH oxidation in tibialis anterior muscle from WT and *ncf1*\* mice (n=5). **d** Exercise signaling in quadriceps muscle from WT and *ncf1*\* mice (n=6 Resting, n=8 Exercise) **e** Total proteins levels of p38 MAPK, ERK 1/2, Total alpha2 AMPK, Total ACC and Coomassie staining as loading control. * denotes P< 0.05 compared to resting condition. For **a** Paired t-test was performed for statistical analysis, * denotes P< 0.05 compared to resting condition. For **c, d** Two-way ANOVA was performed to test for effects of exercise (Exer) genotype (Geno), and interaction (Int), followed by Tukey’s *post hoc* test to correct for multiple comparisons. #, ### denotes P<0.05 and p< 0.001, respectively compared to the WT group. Individual values and mean ± SEM are shown. Scale Bar= 50 μm

### The exercise-induced pro-oxidative shift measured by DCFH requires NOX2 activity in mice

To dissect the ROS source during exercise, we used a previously described p47phox-mutated mice (*ncf1*\*) harboring a loss-of-function mutation in the regulatory NOX2 subunit, p47phox^20^. An acute moderate-intensity treadmill exercise bout (65% maximal running speed) for 20 min increased DCFH oxidation (+ 45%) in WT mice, a response that was completely abolished in p47phox-deficient *ncf1*\* mice (Fig. 1C). This observation was found to be independent of alterations in antioxidant enzyme abundance in TA muscles from *ncf1*\* mice compared to WT mice (Supplementary Fig. 1B). This shows that NOX2-activity is required for moderate-intensity exercise-induced DCFH oxidation in mice.

### Exercise-stimulated p38 MAPK^Thr180/Tyr182^ phosphorylation is reduced in *ncf1*\* quadriceps muscle

Exercise-induced ROS production has been suggested to activate a number of kinases linked to glucose uptake-regulation in skeletal muscle^21^. Interestingly, p-p38 MAPK^Thr180/Tyr182^ levels were lower in quadriceps muscle of *ncf1*\* mice compared to WT mice after exercise (Fig. 1D), but not in TA (Supplementary Fig. 1C) or soleus muscles (Supplementary Fig. 1D). The phosphorylation of ERK^Thr202/Tyr204^, AMPK^Thr172^, and its substrate ACC^Ser212^ did not differ significantly between genotypes (Fig. 1D and Supplementary Fig.1C+D). We also measured SERCA, eEF2^Thr57^, CaMKII^Thr287^ and TBC1D1^Ser231^ but found no genotype-difference (data not shown). No genotype-difference was observed for total AMPKα2, p38 MAPK, ERK 1/2 or ACC in any of the studied muscles (Fig. 1E and Supplementary Fig. 2). The variable responsiveness of these kinases supports that the mice performed moderate-intensity exercise and shows that NOX2 is not consistently required for activation of these kinases.

### NOX2 is required for increased cytosolic ROS during exercise

Since DCFH oxidation does not provide information about the ROS source, we further investigated the subcellular redox changes in response to exercise in skeletal muscle using a recently described redox histology method^18^ (see graphical depiction in Supplementary Fig. 3A). This method enables the preservation and visualization of the redox state of a transfected redox-sensitive GFP 2 (roGFP2)-Orp1, targeted to cytosolic and mitochondrial compartments following *in vivo* exercise. The Orp1 domain facilitates roGFP2 oxidation in the presence of H_2_O_2_, to elicit a ratiometric change, with an increase in 405 nm and a decrease in 470 nm fluorescence. As such, the ratio between the two wavelengths is a measure of H_2_O_2_ production in the targeted compartment, here the cytosol and the mitochondria^22^.

Mito-roGFP2-Orp1 was located in the mitochondrial compartment (co-localizing with tetramethylrhodamine, ethyl ester, Supplementary Fig. 3B-C) and shown to be sensitive to H_2_O_2_ (Supplementary Fig. 3D). Interestingly, oxidation of Mito-roGFP2-Orp1 probe was lowered similarly in both genotypes by exercise (Fig. 2A). In contrast, cytosolic roGFP2-Orp1 oxidation showed a main effect of genotype (Fig. 2B), driven by an exercise-induced increase in roGFP2 oxidation in WT but not *ncf1*\* mice.

**Figure 2.**
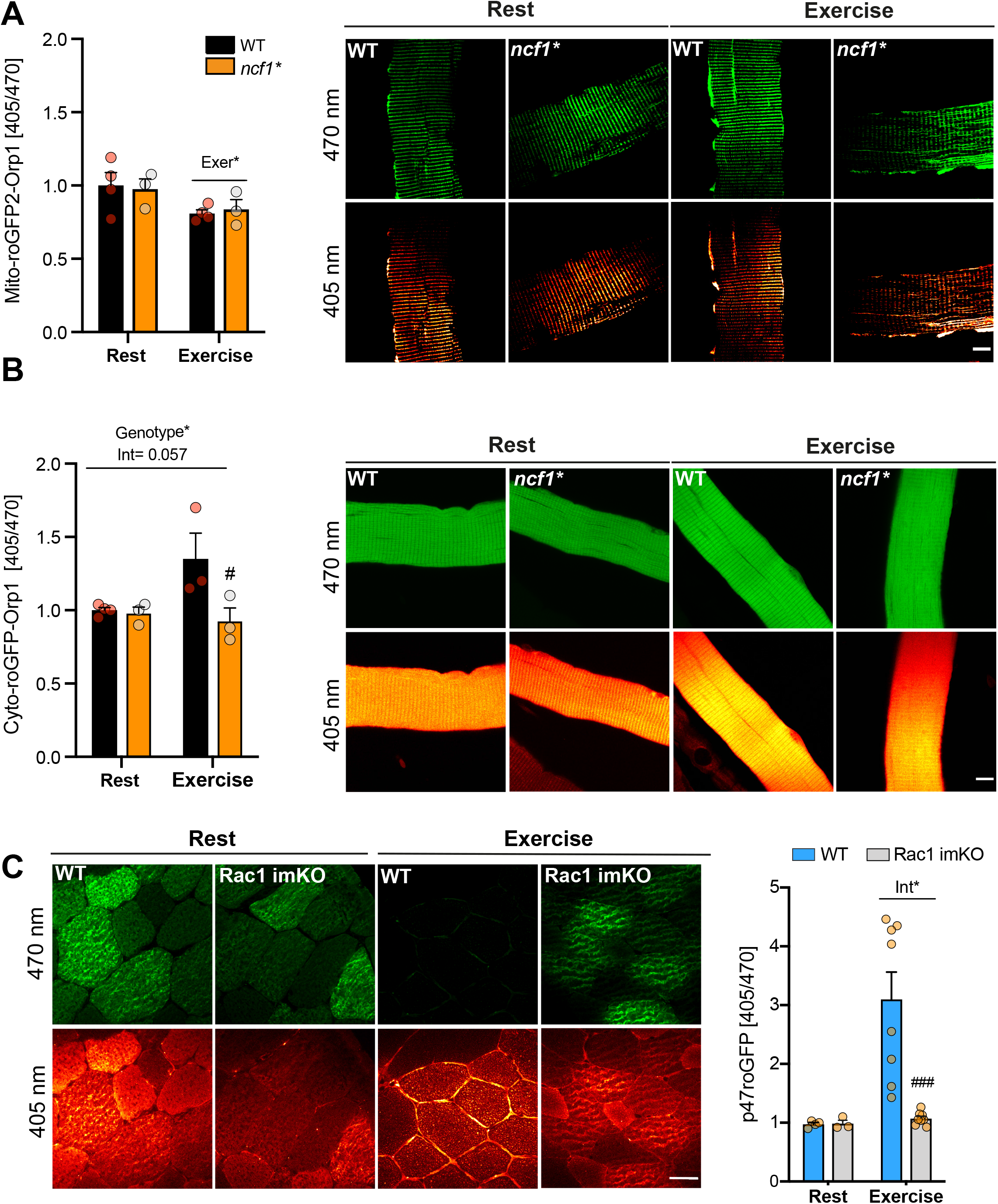
NOX2 is a major ROS source during exercise in skeletal muscle. Subcellularly-targeted redox sensitive GFP2 (roGFP2) were electroporated in *ncf1*\* and p47roGFP in inducible muscle-specific Rac1 mice. Representative image and quantification of **a** Mito-roGFP2-orp1 (WT, n=4 and *ncf1*\*= 3), **b** cyto-roGFP2-Orp1 in Flexor Digitorum Brevis fibers (WT, n=3-4 and *ncf1*\*= 3) and **c** p47roGFP oxidation in tibialis anterior muscle in WT and Rac1 imKO mice (WT, n= 4/8 for rest/exercise, Rac1 imKO, n= 3/8 8 for rest/exercise). Two-way ANOVA was performed to test for effects of exercise (Exer) genotype (Geno), and interaction (Int), followed by Tukey’s *post hoc* test to correct for multiple comparisons. #, ### denotes P<0.05 and p< 0.001, respectively compared to the WT group. Individual values and mean ± SEM are shown. Scale Bar= 10 um (A and B)/ 30 μm (C)

To substantiate that NOX2 activity was required for exercise-stimulated ROS production, we electroporated a biosensor designed to measure NOX2 activity, the p47roGFP construct, into muscles from inducible muscle-specific Rac1 knockout mice (Rac1 imKO), which are predicted to lack functional NOX2 complex. Treadmill exercise caused an acute increase in p47roGFP oxidation in WT TA muscle which was completely absent in Rac1 KO mice, showing that Rac1 is essential for NOX2 activation in skeletal muscle (Fig. 2C). A similar dependence of NOX2 activation on Rac1 was observed in electrically stimulated FDB fibers *in vitro* (Supplementary Fig. 3H). The absent p47roGFP oxidation in Rac1 imKO muscles was not explicable by differences in antioxidant enzyme abundance (Supplementary Fig. 3I). In accordance, exercise-stimulated DCF oxidation was completely absent in Rac1 imKO compared to WT littermates (Fig. S2G) with no differences under resting conditions (Supplementary Fig. 2I).

Collectively, these results show that NOX2 is activated and constitutes a major source of cytosolic ROS production during endurance-type exercise in skeletal muscle. In contrast, mitochondrial ROS production is lowered during acute exercise independently of NOX2.

### NOX2 is required for exercise-stimulated glucose uptake

Given that Rac1 imKO mice display a severe reduction in treadmill exercise-stimulated glucose uptake and GLUT4 translocation^17^, we reasoned that if Rac1 was acting via NOX2 then *ncf1*\* mice should show a similar reduction. We first conducted a general characterization of *ncf1*\* mice. Similar total body weight but reduced fat mass was observed in *ncf1*\* mice compared to age-matched WT mice (Supplementary Fig. 4). Respiratory exchange ratio (RER) was lower in *ncf1*\* mice during the light phase compared to WT mice (Fig. 3A), despite a similar habitual activity observed between genotypes (Fig. 3B). Thus, *ncf1*\* mice, similar to Rac1 imKO mice^23^, display a shift towards fat oxidation at rest and during fasting.

**Figure 3.**
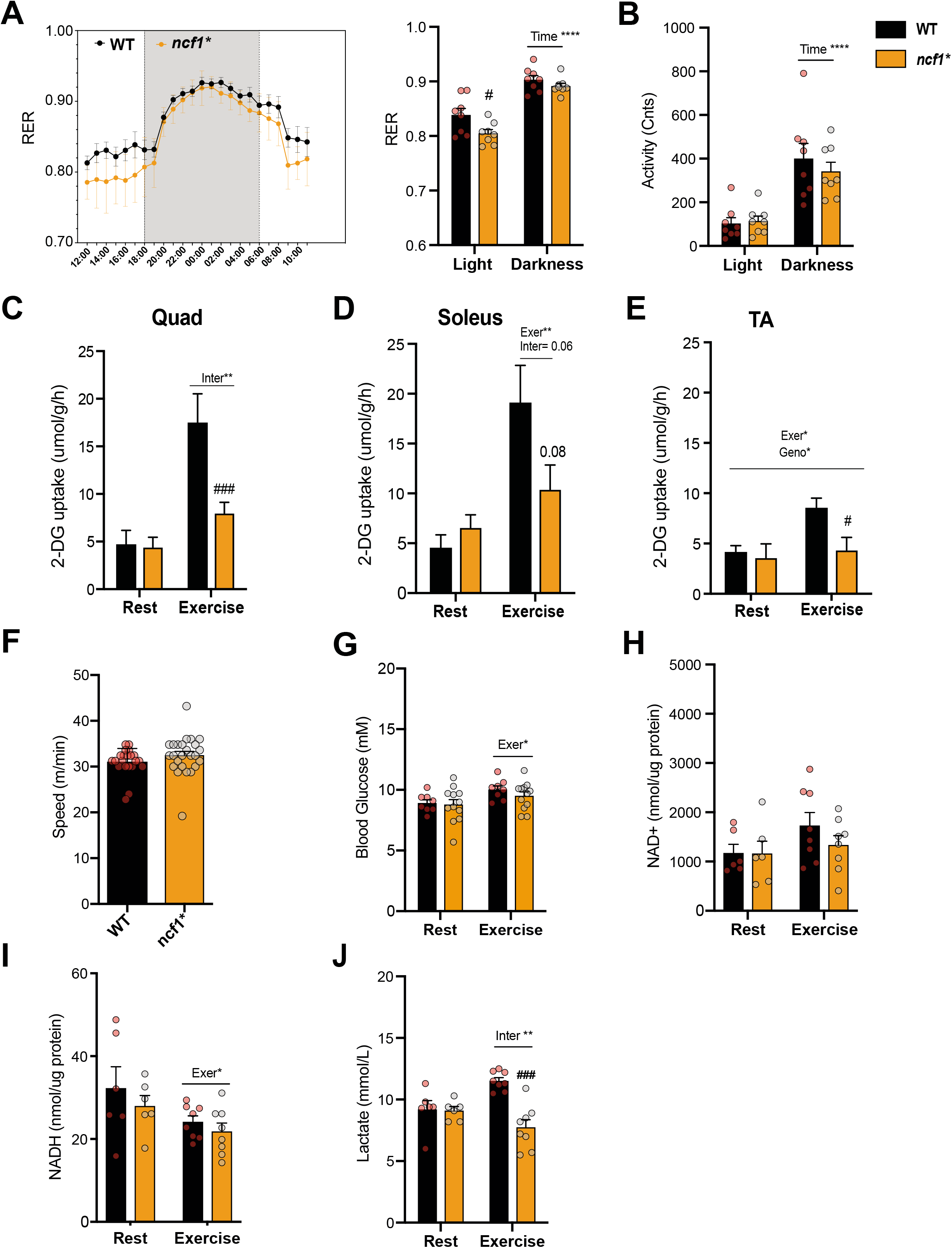
Exercise-stimulated glucose uptake is reduced in *ncf1*\* mice. Different parameters were compared between p47phox mutated mice (*ncf1*\*) and wild-type (WT) mice. **a** Respiratory exchange ratio (RER) during light and dark cycle and **b** habitual activity (n=8 per group). **c-e** *In vivo* running-induced (20 min, 65% maximal running speed) 2-deoxy-glucose uptake (2DG) uptake in quadriceps (Quad) soleus and tibialis anterior (TA) muscles from WT and *ncf1*\* mice (n=12-16). **f** Maximal running speed (n=15 per group), **g** blood glucose concentration after rest/exercise (n=8-12), **h** NAD+, **i** NADPH levels in quadriceps muscle (n=6-8 for rest/exercise), and **j** Plasma lactate concentration (n=6-8 for rest/exercise). Unpaired t-test (**f**) and two-way ANOVA were performed to test for effects of exercise (Exer) genotype (Geno), and interaction (Int), followed by a Tukey’s *post hoc* test to correct for multiple comparisons. #, ### denotes P< 0.05 and p< 0.001, respectively compared to the WT group. Individual values and mean ± SEM are shown.

Next, we determined whether NOX2-dependent ROS generation is required for exercise-stimulated glucose uptake. Each mouse was exercised at a relative work load corresponding to 65% of its maximum running capacity for 20 min. Exercise-induced 2- [^3^H] deoxyglucose (2DG) uptake was markedly attenuated in quadriceps, soleus, and TA muscles from *ncf1*\* mice compared to WT (Fig. 3C-E). Importantly, the reduction of exercise-stimulated glucose uptake in *ncf1*\* mice was not due to differences in maximal running capacity (Fig. 3F), blood glucose levels (Fig. 3G) or intramuscular energetics evaluated as NAD+ and NADH levels (Fig. 3H-I). However, plasma lactate levels were increased by exercise in WT but not in *ncf1*\* *mice* (Fig. 3J), consistent with a reduced reliance on glycolysis for energy production. No differences were observed in muscle fiber size (Fig. 4A-C), fiber-type composition (Fig. 4D+E), mitochondrial complex expression (Fig. 4F+G, quantifications in Supplementary Fig. 5) or capillary density (Fig. 4H). Thus, *ncf1*\* in comparison to WT mice were markedly impaired in their ability to stimulate glucose uptake *in vivo* across multiple muscles, without indications of altered glucose delivery or oxidation capacity.

**Figure 4.**
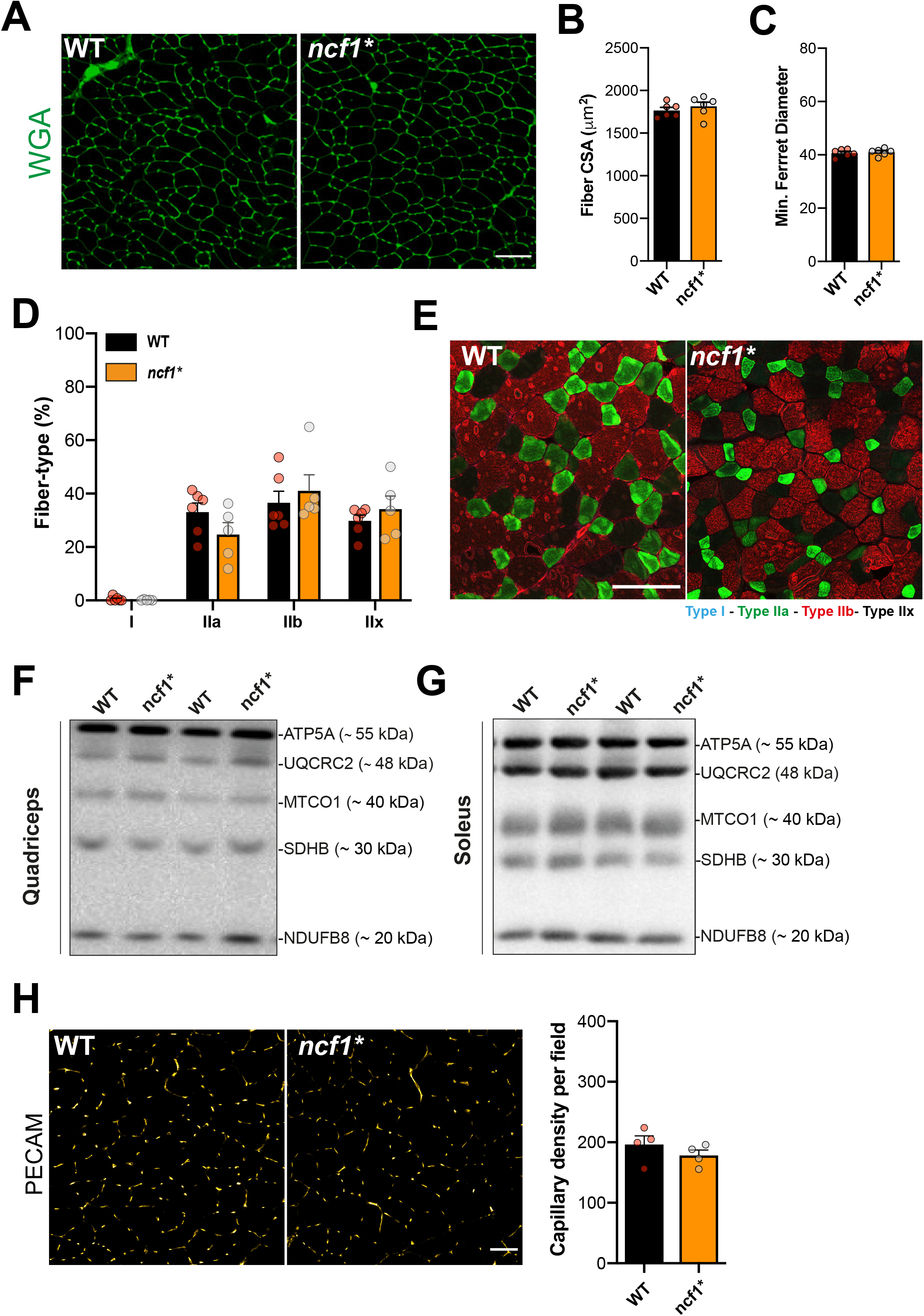
*Ncf1*\* mice show similar muscle size, fiber-type composition, mitochondrial content and capillary density compared to WT group. **a** Fiber-size parameters in WT and *ncf1*\* mice were determined using Wheat Germ Agglutinin (WGA) staining. **b** Myofiber Cross sectional area, and **c** minimum (min.) Ferret diameter. **d** Quantification and **e** representative merged image of fiber-type staining where Type I fibers (blue), Type IIa (green), Type IIb (red) Type IIx (non-stained) in WT (n= 6) and *ncf1*\* (n= 5) TA muscles. Subunits of mitochondrial oxidative phosphorylation complexes were determined in **f** quadriceps and **g** soleus muscle lysates (n=14 per group). **h** Capillary density was estimated by PECAM immunostaining in tibialis anterior sections. Two-way ANOVAs was performed to test for effects of exercise (Exer) genotype (Geno), and interaction (Int), followed by Tukey’s *post hoc* test to correct for multiple comparisons. Individual values and mean ± SEM are shown. Scale Bar= 100 μm

### NOX2 is required for exercise-stimulated GLUT4 translocation

Next, we tested if the reduction in exercise-stimulated glucose uptake in *ncf1*\* mice could be explained by reduced GLUT4 translocation as observed in the Rac1 imKO^17^. Consistent with this notion, the exercise-stimulated increase in surface-membrane GLUT4-GFP-*myc* in TA muscle from WT was virtually abolished in *ncf1*\* mice (Fig. 5A). Importantly, total protein abundance of GLUT4 and HKII (Fig. 5B-D, quantification in Supplementary Fig. S6). Taken together, these data demonstrate that *ncf1*\* mice phenocopy the Rac1 imKO mice in terms of pronounced impairments in exercise-stimulated muscle glucose uptake and GLUT4 translocation^17^ This strongly suggests that the common denominator between p47phox and Rac1, NOX2 activation, is required for exercise-stimulated glucose uptake and GLUT4 translocation.

**Figure 5.**
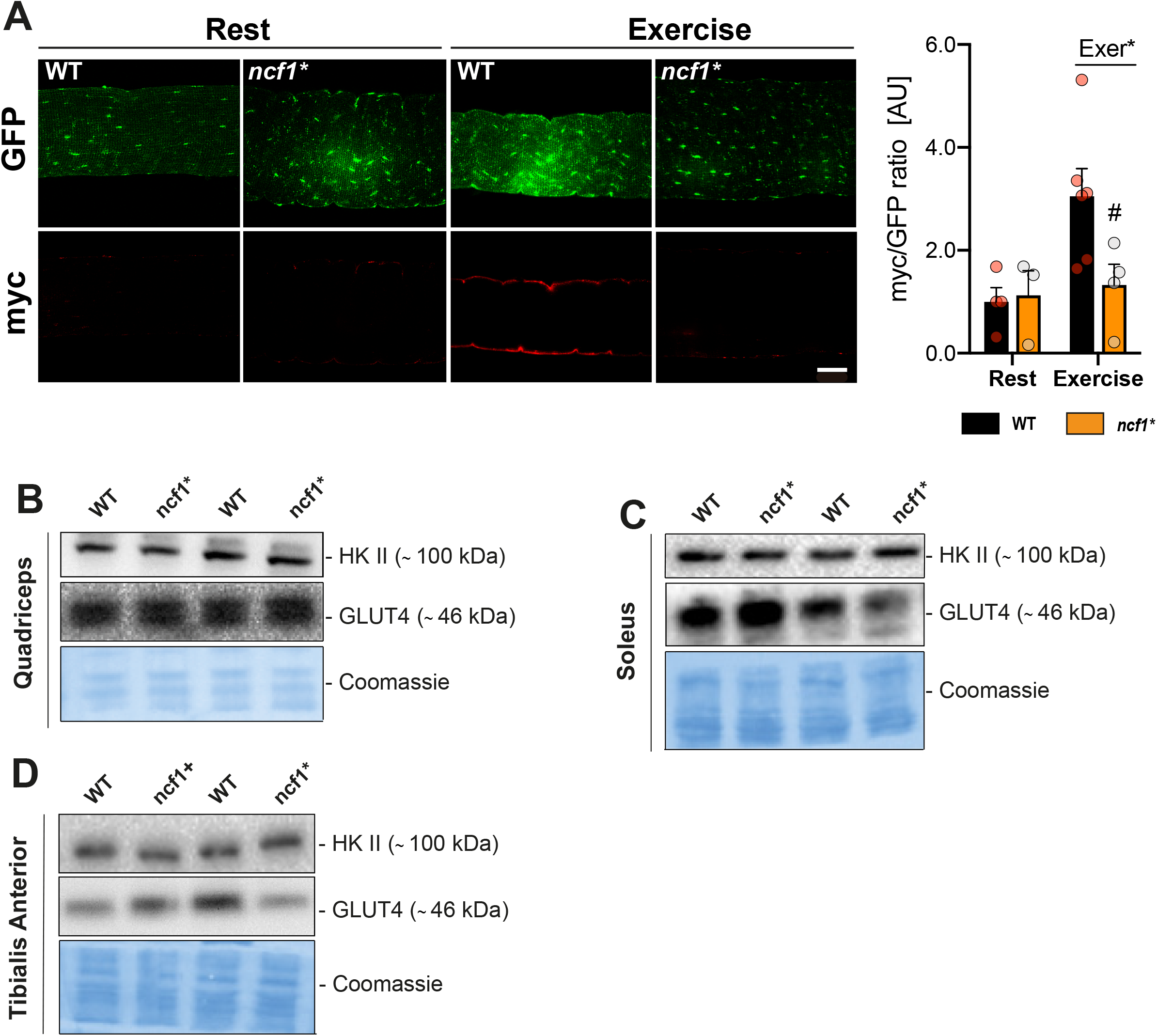
Exercise-stimulated GLUT4 translocation requires NOX2 activity. **a** The GLUT4-myc-GFP construct was electroporated into tibialis anterior muscles of both WT and *ncf1*\* mice. Non-permeabilized muscle fibers from exercised (20 min, 65% maximal running speed) or resting mice were stained with anti-*myc* antibody and imaged by confocal microscopy (n=4). Total endogenous GLUT4 and HK II were determined by western blot in the following muscles **b** quadriceps, **c** soleus, and **d** TA muscles in WT and *ncf1*\* mice (n=14). Unpaired t-test (**b-d**) and two-way ANOVA were performed to test for effects of exercise (Exer) genotype (Geno), and interaction (Int), followed by a Tukey’s *post hoc* test to correct for multiple comparisons. Individual values and mean ± SEM are shown. Scale bar= 20 μm

## DISCUSSION

The source of ROS during exercise and the role of ROS in the acute and chronic adaptations to exercise, in particular in the context of non-damaging moderate-intensity *in vivo* exercise, has remained undetermined for more than 30 years^24^ The current study showed that moderate-intensity endurance exercise acutely caused a pro-oxidative shift in both humans and mice. Strikingly, our results demonstrated that NOX2 was required for this shift and was a major cytosolic ROS source during moderate-intensity exercise. Furthermore, NOX2 activity was necessary for exercise-stimulated GLUT4 translocation and glucose uptake in skeletal muscle, providing a novel molecular explanation of why Rac1 imKO mice have a strong impairment of exercise-stimulated GLUT4 translocation and glucose uptake.

The myocellular ROS source(s) during exercise conditions has long been debated, with both cytosolic and mitochondrial sources having been proposed based on pharmacological *in vitro/ex vivo* studies^3, 25, 26, 27, 28^. Presently, we observed an increase in oxidants measured by DCFH oxidation in exercising mouse and human muscle. Remarkably, despite the non-specific nature of DCF fluorescence, we found that the increase in exercise-stimulated DCFH oxidation was virtually abolished in mice lacking either Rac1 or p47phox components of NOX2. Then, we used genetically-encoded redox probes targeted to different myocellular compartments to study the ROS sources during *in vivo* exercise. Exercise promoted an oxidation of cytosol-targeted roGFP2-Orp1 biosensor in WT but not in NOX2 deficient mice. On the other hand, mitochondria-targeted roGFP2-Orp1 probe oxidation was reduced by exercise independently of NOX2. The NOX2 complex-targeted p47roGFP biosensor oxidation was increased by exercise in WT muscles but not in Rac1 deficient mice. Overall, this demonstrates that a functional NOX2 complex, but not mitochondria, is required for cytosolic ROS production induced by exercise in concordance with previous *in vitro* studies.

We and others have shown that the small rho family GTPase Rac1 is necessary for insulin, contraction and passive stretch-induced skeletal muscle glucose uptake^13, 14, 29^. We have previously found this Rac1-dependency of glucose uptake and GLUT4 translocation to be more pronounced during *in vivo* treadmill exercise compared to *in vitro* contractions^17^ The GLUT4 trafficking field has mostly focused on Rac1 as an orchestrator of the dynamic reorganization of the cortical actin cytoskeleton consisting of the cytoplasmic β and γ actin-isoforms^15^. This actin-reorganization was shown to be crucial for insulin-stimulated GLUT4 translocation in muscle cells^30^. However, the cytoplasmic β and γ actin-isoforms are both downregulated during muscle differentiation and whether dynamic reorganization occurs in adult skeletal muscle fibers similarly to in cells is presently unclear^31^. Our present data suggest that NOX2 is a vital downstream mediator of the effect of Rac1 on exercise-induced glucose uptake and GLUT4 translocation since the Rac1 imKO phenotype is shared by mice lacking another essential regulatory subunit of NOX2, p47phox. In addition, our *in vivo* glucose uptake data provide a likely physiological mechanism explaining why exogenous antioxidants inhibit *ex vivo* contraction- and stretch-stimulated glucose uptake^10, 11^. The relative dependence on actin and NOX2 and whether NOX2 and actin interact in muscle to regulate glucose uptake, as suggested in other cell types^32^, should be clarified in future studies.

In the current study, the exercise-stimulated phosphorylation of p38MAPK was reduced in *ncf1*\* mice compared to WT, in particular in quadriceps muscle. As p38 MAPK has been proposed as a regulator of muscular glucose uptake^11^, we cannot exclude that reduced p38 MAPK contributed to the reduced glucose uptake we observed in *ncf1*\* mice. However, the reduction in exercise-stimulated kinase signaling is less consistent than the reduction in glucose uptake in the different muscles analyzed in the *ncf1*\* mice. Moreover, Rac1 imKO mice share the same glucose uptake phenotype without showing reductions in stretch-stimulated signaling, including p38 MAPK^17^ Instead, we speculate that the shared reduction in exercise-stimulated glucose uptake in *ncf1*\* and Rac1 imKO mice might relate to a shared and yet undetermined redox-sensitive signaling mechanism. Worth noting, we believe NOX2 to signal independently of the AMPK-TBC1D1 signaling axis, based on the previous work by us and others^13, 33, 34, 35, 36^. In conjunction with the current study, strongly suggesting that Rac1 regulates glucose uptake via NOX2, we propose that NOX2 also signals independently of AMPK to regulate glucose uptake during moderate-intensity exercise. Future studies should work to identify the shared cell signaling traits between different NOX2-deficient mouse models using e.g. redox proteomics^37^.

In conclusion, this study showed for the first time that NOX2 is activated during moderate-intensity exercise in human and mouse skeletal muscle and is the primary source of ROS under such conditions. Furthermore, our comparison of two mouse models lacking regulatory NOX2 subunits showed that lack of ROS generation during exercise strongly impaired muscle glucose uptake and GLUT4 translocation during exercise. This indicates that NOX2 is a major source of ROS generation during exercise and that NOX2-dependent ROS production is an important signal for increasing muscle glucose uptake during exercise.

## MATERIAL AND METHODS

### Animals

All experiments were approved by the Danish Animal Experimental Inspectorate (2015–15–0201–00477). Inducible muscle-specific male Rac1 mice (imKO) and littermate control mice were generated by crossbreeding Rac1^fl/fl^ mice^38^ with mice expressing Cre recombinase from a tetracycline-controlled transactivator coupled to the human skeletal actin promoter^39^. For muscle-specific Rac1 deletion, all mice were treated with doxycycline in the drinking water around 10-14 weeks of age (1g/L; Sigma Aldrich) for 3 weeks followed by a wash out period of 3 weeks.

Male B10.Q wild-type (WT) and B10.Q. p47phox mutated (*ncf1*\*) mice contain a point mutation in exon 8^20^. All mice were group-housed maintained on a 12:12-h light-dark cycle, at 21° C, and received standard rodent chow diet (Altromin no. 1324; Chr. Pedersen, Denmark) and water ad libitum.

### Human experiments

Three young (Age 29 ± 3.56y BMI 24.8 ± 1.76 kg/m^2^), healthy men gave their written, informed consent to participate in the study approved by the Regional Ethics Committee for Copenhagen (H-16040740) and complied with the ethical guidelines of the Declaration of Helsinki II. The volunteers visited the laboratory on two separate days. The first day, the subjects completed an incremental test on a Monark Ergomedic 839E cycle ergometer (Monark, Sweden) to determine peak power output (PPO). On the second day, each subject performed a 30-min exercise trial at the intensity corresponding to ~65% of their individual peak power output in fasted state. Muscle biopsies were obtained from the m. vastus lateralis under local anesthesia [~3ml xylocaine (20mg ml–1lidocaine), Astra, Stockholm, Sweden] before and after exercise using a 5 mm Bergstrom needle with suction. The muscle biopsies were embedded in optimal cutting temperature compound (tissue-tek) and frozen in liquid nitrogen-cooled isopentane and stored at −80°C for further analysis.

### In vivo gene transfer in adult skeletal muscle

For the *in vivo* transfection experiments, mice were anesthetized with 2-3% isoflurane. Hyaluronidase (Sigma-Aldrich) dissolved in sterile saline solution (0.36 mg/ml) was injected subcutaneously on the plantar side of the foot near but not into the FDB and intramuscularly in TA muscle, followed by 20 and 40 μg plasmid injections 1 hour later, respectively. FDB muscle electroporation was performed by delivering 10 electrical pulses at an intensity of 75 V/cm, 10-ms pulse duration, 200-ms pulse interval using acupuncture needles (0.20 × 25 mm, Tai Chi, Lhasa OMS) connected to an ECM 830 BTX electroporator (BTX Harvard Apparatus). TA electroporation was performed following the same protocol but raising the intensity to 100 V/cm (#45-0101 Caliper Electrode, BTX Caliper Electrodes, USA). The p47-roGFP construct used to determine NOX2 activity was a kind gift from Professor George G. Rodney^40^. The Cyto-roGFP2-Orp1 (Addgene #64991) and Mito-roGFP2-Orp1 (Addgene #64992) plasmids were gifts from Professor Tobias Dick. The GLUT4-myc-GFP construct was a gift from Professor Jonathan Bogan (Addgene #52872). After electroporation, mice were allocated to their cages for 10 days before performing any experiments.

### Metabolic Chambers

O_2_ uptake (VO_2_) and CO_2_ production (VCO_2_) were measured using a CaloSys apparatus (TSE Systems, Bad Homburg, Germany). The data presented is the average for light and dark periods measured over 2 consecutive days in WT and *ncf1*\* mice. The respiratory exchange ratio (RER) was calculated as VCO_2_ production/VO_2_ uptake.

### Maximal running tests

Mice were acclimated to the treadmill three times (15 min at 0.16 m/s) (Treadmill TSE Systems) a week prior to the maximal running tests. The maximal running test started at 0.16 m/s for 300 s with 10 % incline, followed by a continuing increase (0.2 m/s) in running speed every 60s until exhaustion. Exhaustion was defined as the point at which instead of running on the treadmill, the mice fall back on the grid 3 times within a 30 sec period. Maximal running speed was determined as the last completed stage during the incremental test.

### FDB fiber isolation

Muscle fibers were obtained by enzyme digestion of the whole muscle with collagenase type II (1.5 mg/ml) (Worthington, Lakewood, NJ) for 90 min at 37°C followed by mechanical dissociation with fire polished Pasteur pipettes. Isolated fibers were seeded in ECM Gel-coated (Sigma-Aldrich) cell culture dishes in DMEM supplemented with 10% fetal bovine serum. After 16 h of seeding, the fibers were used for experimentation.

### Live imaging

FDB fibers were imaged in phenol red-free growth medium (5 % horse serum) while kept at 95 % O_2_ and 5 % CO_2_ and 37°C using a Pecon Lab-Tek S1 heat stage. Electrical stimulation protocol (10 V, 1 Hz, 0.1 ms duration) was delivered to isolated FDBs for 15 min via field electric field stimulation device (RC-37FS, Warner instruments, USA). Fibers were loaded with 20 nM tetramethylrhodamine, ethyl ester (TMRE^+^, Life Technologies) for 30 min before imaging.

### Redox histology

TA muscles were dissected and embedded in optimum cutting temperature (OCT) medium from Tissue-Tek, frozen in melting isopentane and kept at −80 °C until processing. An independent set of TA muscles were used for western blotting. Total oxidant levels were estimated as previously described^12^. Briefly, 10 μm thickness muscle cryosections were incubated with 5 μM of 2’,7’-dichlorodihydrofluorescein diacetate (DCFH; Molecular Probes, Eugene) and allowed to dry overnight at room temperature in dark. Muscle membranes were visualized using Texas red-labeled wheat germ agglutinin (WGA; Molecular Probes). The redox histology was performed as previously described^18^. Whole FDB muscles were immediately immersed in PBS containing freshly dissolved NEM (100 mM) in ice and then fixed in 4% PFA in PBS (pH 7.4) plus 100 mM NEM for 2 hours at 4° C. p47roGFP-transfected TA cryosections were incubated with PBS containing 50 mM N-Ethylmaleimide (NEM) for 10 min at 4°C, followed by fixation using 4% paraformaldehyde for 10 min at room temperature. All samples were mounted in mounting medium (Vectashield, USA).

### Cryosection Immunostaining

Cryosections of TA muscle (cut in transverse orientation) were stained with monoclonal anti-myosin heavy chain (MyHC) antibodies (DSHB, University of Iowa): BA-D5 (IgG2b, supernatant, 1:100 dilution) specific for MyHC-I, SC-71 (IgG1, supernatant, 1:100 dilution) specific for MyHC-2A, and BF-F3 (IgM, purified antibody, 1:100 dilution) specific for My-HC-2B. Type 2X fibers are not recognized by these antibodies, and so appear black. Three different secondary antibodies (Life Technologies, Carlsbad, USA) were used to selectively bind to each primary antibody: goat-anti-mouse IgG2b, conjugated with Alexa 647 fluorophore (to bind to BA-D5); goat-anti-mouse IgG1, conjugated with Alexa 488 fluorophore (to bind to SC-71); goat-anti-mouse IgM, conjugated to Alexa 555 fluorophore (to bind to BF-F3). Muscle sections, 10 μm thick, were fixed in 2 % PFA for 30 min and permeabilized with 0.1 % Triton-X100 for 10 min. After washing in immunobuffer (IB = 0.25 % BSA, 50mM glycine, 0.033 % saponin and 0.05 % sodium azide diluted in PBS), sections were blocked for 1 h in 2 % BSA, then briefly washed twice with IB for 5 min. A solution with all the primary antibodies diluted in IB was then prepared, and sections were incubated for 2 h at 37 °C. After 3 washes (10 min each) with IB, sections were incubated for 1 h at 37 °C with a solution containing the three different secondary antibodies diluted in IB. After 3 washes with IB (10 min each) and a brief rinse in PBS, sections were mounted with Fluoromount (Sigma-Aldrich, St. Louis, USA). Capillary density was evaluated using a PECAM1 immunostaining in TA cryosections. After fixation and blocking steps as described above, anti-PECAM1 antibody (SCBT, M-20) diluted 1:100 in 1% BSA was incubated overnight at 4° C. After 3 washes (5 min each), an anti-goat Alexa 488-conjugated secondary antibody was incubated for 1h. After 3 washes with IB (5 min each) sections were mounted.

### Imaging and image analysis

For all live-imaging experiments confocal images were collected using a 63x 1.4 NA oil immersion objective lens on an LSM 780 confocal microscope (Zeiss) driven by Zen 2011. TMRE^+^ fluorescence was detected using the excitation-emission λ545–580/590 nm. For the roGFP biosensors images, raw data of the λ405- and 488-nm laser lines were exported to ImageJ as 16-bit TIFFs for further analysis. Data are presented as normalized fluorescence ratio (λ 405/488 nm) normalized to the resting WT group. For fiber-type immunostainings, images were collected using a dry 20x 0.8 NA Plan Apo objective on an LSM 710 confocal microscope (Zeiss) driven by Zen 2012. The used excitation laser lines were 488, 561 and 633 nm, respectively, assembled by Zeiss. Three tracks were sequentially used for acquisition, with 488 and 633 channels recorded with PMTs, while the 561 channel was recorded with a GaAsP detector array. The matching dichroic mirrors were used for all channels, and the pinhole was set at 1 AU for 580 nm. Images were exported to ImageJ and the composition of different MHC fiber types were quantified by counting sections of each muscle bed (Type I, blue; Type IIa, green; Type IIb, red; Type IIx, black) using the Cell Counter plugin.

### Running-stimulated muscle 2-deoxyglucose (DG) uptake

Muscle-specific 2DG uptake was measured as previously described^33^. [^3^H]2DG (PerkinElmer) was injected intraperitoneally (as a bolus in saline, 10 μL/g body weight) containing 0.1 mmol/L 2DG and 50 μCi/mL [^3^H]2DG corresponding to ~12 μCi/mouse) into fed mice immediately before the exercise bout. Blood glucose was measured before and immediately after 20 min of exercise. Mice were euthanized by cervical dislocation, and muscle tissues and plasma samples quickly frozen in liquid nitrogen and stored at −80°C until analysis. Samples were deproteinated using 0.1 mol/L Ba (OH)_2_ and 0.1 mol/L ZnSO_4_. The total muscle [^3^H]2DG tracer activity found in 2DG-6-phosphate was divided by the area under the curve of the specific activity at time points 0 and 20 min multiplied with the average blood glucose at time points 0 and 20 min. This was related to muscle weight and the time to obtain the tissue-specific 2DG uptake as micromoles per gram per hour.

### Exercise-stimulated GLUT4 translocation

GLUT4 translocation was measured as recently described^41^. Briefly, rapidly dissected TA muscles were fixed by immersion in ice-cold 4% in paraformaldehyde in PBS for 4 h. Individual fibers were teased from fixed muscle with fine forceps under a dissection microscope. Non-permeabilized isolated muscle fibers were incubated in 1% BSA in PBS for 1 h min and then incubated with an anti-myc antibody (rabbit polyclonal) overnight (CST, #2278). The next day fibers were incubated with a secondary antibody conjugated with Alexa Fluor 568 (Invitrogen, UK). The muscle fibers were mounted in Vectashield mounting medium.

### Western blot analyses

Tissue was homogenized for 1 min at 30 Hz using a Tissue Lyser in ice cold lysis buffer (0.05 mol/L Tris Base pH 7.4, 0.15 mol/L NaCl, 1 mmol/ L EDTA and EGTA, 0.05mol/L sodium flouride, 5 mmol/L sodium pyrophosphate, 2 mmol/L sodium orthovanadate, 1 mmol/L benzamidine, 0.5% protease inhibitor cocktail (P8340, Sigma Aldrich), and 1% NP-40). After rotating end-over-end for 30 min, lysate supernatants were collected by centrifugation (18,327 g) for 20 min at 4°C. Lysate protein concentrations were determined using BSA standards and bicinchoninic acid assay reagents (Pierce). Total protein and phosphorylation levels of relevant proteins were determined by standard immunoblotting techniques, loading equal amounts of protein. The primary antibodies used p-AMPK^Thr172^ (Cell Signaling Technology (CST)), #2535S), p-p38 MAPK^Thr180/Tyr182^ (CST, #9211), Erk1/2^Thr202/Tyr204^ (CST, #9101), GLUT4 p-ACC2 Ser^212^ (Millipore, 03-303), (ThermoFisher Scientific, PA-23052), Rac1 (BD Biosciences, #610650), NOX2 (Abcam, #Ab129068), Catalase (SCBT, sc-271803), MnSOD (Millipore, 06-984), TRX2 (SCBT, sc-50336), actin (CST, #4973) total p38 MAPK (CST, #9212), alpha2 AMPK (a gift from D. Grahame Hardie, University of Dundee), total ERK 1/2 (CST, #9102), and TBC1D1^ser231^ (Millipore #07-2268), OXPHOS cocktail (Abcam, #ab110413), total TBC1D1 (Abcam, #Ab229504) and Hexokinase II (CST, #2867). ACC protein was detected using horseradish peroxidase-conjugated streptavidin from Dako (P0397), dilution 1:3000. All antibodies were optimized for signal linearity. Membranes were blocked for 1 h at room temperature in TBS-Tween 20 containing either 2% skimmed milk or 2% BSA and incubated overnight at 4°C with primary antibody. The membranes were incubated in the corresponding horseradish peroxidase-conjugated secondary antibody for 1 h at room temperature and washed in TBS-T before visualization of the proteins (ChemiDoc™MP Imaging System, Bio Rad).

### Statistical analyses

Results are shown as individual values or means ± S.E.M. Statistical testing was carried out using *t*-tests or two-way (repeated measures when appropriate) ANOVA as applicable. Tukey’s *post hoc* test was performed for multiple comparisons when ANOVA revealed significant main effects or interactions. Statistical analyses were performed using GraphPad Prism v8.

## Acknowledgements

TEJ was supported by a Novo Nordisk Foundation Excellence project grant (#15182). CHO was supported by a Chilean CONICYT PhD Scholarship. JRK was supported by Danish Diabetes Academy PhD stipends. ZL was supported by Chinese Scholarship Council PhD stipends. LS was supported by a Danish Research Council grant 4004-00233 and Novo Nordisk Foundation Excellence grant NNF18OC0032082. EJ was supported by FONDECYT grant 1151293. EAR was supported by the Danish Council for Independent Research/Science 6108-00203. RH was supported by the Swedish Research Council, the Knut and Alice Wallenberg foundations and the Swedish Cancer Society. We thank Kim Anker Sjøberg for pre-testing the human subjects and Betina Bolmgren for technical assistance with measuring 2DG uptake. Imaging data were collected at the Center for Advanced Bioimaging and the Core Facility for Integrated Microscopy, University of Copenhagen, Denmark.

## Author Contribution

Conceptualization, CHO, EJ and TEJ. Methodology, CHO and TEJ. Investigation, CHO, JRK, ZL, ED, JTT, SHR, RH, EAR and LS. Formal analysis, CHO. Visualization, CHO. Writing – Original draft, CHO and TEJ. Writing – Review and Editing, CHO with input from all authors. Funding Acquisition, TEJ. Supervision, EJ and TEJ.

## Data availability

The authors declare that all data supporting the findings of this study are available within the article and its supplementary information files or from the corresponding author upon reasonable request.

## REFERENCES

1. Egan B, Zierath JR. Exercise metabolism and the molecular regulation of skeletal muscle adaptation. Cell metabolism 17, 162–184 (2013).

2. Powers SK, Jackson MJ. Exercise-induced oxidative stress: cellular mechanisms and impact on muscle force production. Physiol Rev 88, 1243–1276 (2008).

3. Sakellariou GK, et al. Studies of mitochondrial and nonmitochondrial sources implicate nicotinamide adenine dinucleotide phosphate oxidase(s) in the increased skeletal muscle superoxide generation that occurs during contractile activity. Antioxidants & redox signaling 18, 603–621 (2013).

4. Powers SK, Radak Z, Ji LL. Exercise-induced oxidative stress: past, present and future. J Physiol 594, 5081–5092 (2016).

5. Jackson MJ. Recent advances and long-standing problems in detecting oxidative damage and reactive oxygen species in skeletal muscle. J Physiol 594, 5185–5193 (2016).

6. Richter EA, Hargreaves M. Exercise, GLUT4, and skeletal muscle glucose uptake. Physiol Rev 93, 993–1017 (2013).

7. Sylow L, Kleinert M, Richter EA, Jensen TE. Exercise-stimulated glucose uptake – regulation and implications for glycaemic control. Nat Rev Endocrinol 13, 133–148 (2017).

8. Kristiansen S, Hargreaves M, Richter EA. Exercise-induced increase in glucose transport, GLUT-4, and VAMP-2 in plasma membrane from human muscle. Am J Physiol 270, E197–201 (1996).

9. Zisman A, et al. Targeted disruption of the glucose transporter 4 selectively in muscle causes insulin resistance and glucose intolerance. Nat Med 6, 924–928 (2000).

10. Sandstrom ME, et al. Role of reactive oxygen species in contraction-mediated glucose transport in mouse skeletal muscle. J Physiol 575, 251–262 (2006).

11. Chambers MA, Moylan JS, Smith JD, Goodyear LJ, Reid MB. Stretch-stimulated glucose uptake in skeletal muscle is mediated by reactive oxygen species and p38 MAP-kinase. J Physiol 587, 3363–3373 (2009).

12. Merry TL, Steinberg GR, Lynch GS, McConell GK. Skeletal muscle glucose uptake during contraction is regulated by nitric oxide and ROS independently of AMPK. Am J Physiol Endocrinol Metab 298, E577–585 (2010).

13. Sylow L, et al. Rac1 is a novel regulator of contraction-stimulated glucose uptake in skeletal muscle. Diabetes 62, 1139–1151 (2013).

14. Sylow L, Moller LL, Kleinert M, Richter EA, Jensen TE. Stretch-stimulated glucose transport in skeletal muscle is regulated by Rac1. J Physiol 593, 645–656 (2015).

15. Chiu TT, Jensen TE, Sylow L, Richter EA, Klip A. Rac1 signalling towards GLUT4/glucose uptake in skeletal muscle. Cell Signal 23, 1546–1554 (2011).

16. Bedard K, Krause KH. The NOX family of ROS-generating NADPH oxidases: physiology and pathophysiology. Physiol Rev 87, 245–313 (2007).

17. Sylow L, et al. Rac1 governs exercise-stimulated glucose uptake in skeletal muscle through regulation of GLUT4 translocation in mice. J Physiol 594, 4997–5008 (2016).

18. Fujikawa Y, et al. Mouse redox histology using genetically encoded probes. Science signaling 9, rs1 (2016).

19. Margaritelis NV, et al. Adaptations to endurance training depend on exercise-induced oxidative stress: exploiting redox interindividual variability. Acta physiologica (Oxford, England) 222, (2018).

20. Hultqvist M, Olofsson P, Holmberg J, Backstrom BT, Tordsson J, Holmdahl R. Enhanced autoimmunity, arthritis, and encephalomyelitis in mice with a reduced oxidative burst due to a mutation in the Ncf1 gene. Proceedings of the National Academy of Sciences of the United States of America 101, 12646–12651 (2004).

21. He F, Li J, Liu Z, Chuang CC, Yang W, Zuo L. Redox Mechanism of Reactive Oxygen Species in Exercise. Front Physiol 7, 486 (2016).

22. Schwarzlander M, Dick TP, Meyer AJ, Morgan B. Dissecting Redox Biology Using Fluorescent Protein Sensors. Antioxidants & redox signaling 24, 680–712 (2016).

23. Sylow L, et al. Rac1 signaling is required for insulin-stimulated glucose uptake and is dysregulated in insulin-resistant murine and human skeletal muscle. Diabetes 62, 1865–1875 (2013).

24. Davies KJ, Quintanilha AT, Brooks GA, Packer L. Free radicals and tissue damage produced by exercise. Biochem Biophys Res Commun 107, 1198–1205 (1982).

25. Trewin AJ, et al. Acute HIIE elicits similar changes in human skeletal muscle mitochondrial H2O2 release, respiration, and cell signaling as endurance exercise even with less work. Am J Physiol Regul Integr Comp Physiol 315, R1003–R1016 (2018).

26. Goncalves RL, Quinlan CL, Perevoshchikova IV, Hey-Mogensen M, Brand MD. Sites of superoxide and hydrogen peroxide production by muscle mitochondria assessed ex vivo under conditions mimicking rest and exercise. The Journal of biological chemistry 290, 209–227 (2015).

27. Wong HS, Dighe PA, Mezera V, Monternier PA, Brand MD. Production of superoxide and hydrogen peroxide from specific mitochondrial sites under different bioenergetic conditions. The Journal of biological chemistry 292, 16804–16809 (2017).

28. Henriquez-Olguin C, et al. NOX2 Inhibition Impairs Early Muscle Gene Expression Induced by a Single Exercise Bout. Front Physiol 7, 282 (2016).

29. Ueda S, et al. Crucial role of the small GTPase Rac1 in insulin-stimulated translocation of glucose transporter 4 to the mouse skeletal muscle sarcolemma. FASEB journal: official publication of the Federation of American Societies for Experimental Biology 24, 2254–2261 (2010).

30. Chiu TT, Patel N, Shaw AE, Bamburg JR, Klip A. Arp2/3- and cofilin-coordinated actin dynamics is required for insulin-mediated GLUT4 translocation to the surface of muscle cells. Mol Biol Cell 21, 3529–3539 (2010).

31. Madsen AB, et al. beta-actin shows limited mobility and is only required for supraphysiological insulin-stimulated glucose transport in young adult soleus muscle. Am J Physiol Endocrinol Metab, (2018).

32. Wilson C, et al. A Feed-Forward Mechanism Involving the NOX Complex and RyR-Mediated Ca2+ Release During Axonal Specification. J Neurosci 36, 11107–11119 (2016).

33. Sylow L, et al. Rac1 and AMPK Account for the Majority of Muscle Glucose Uptake Stimulated by Ex Vivo Contraction but Not In Vivo Exercise. Diabetes 66, 1548–1559 (2017).

34. Jensen TE, et al. Contraction-stimulated glucose transport in muscle is controlled by AMPK and mechanical stress but not sarcoplasmatic reticulum Ca(2+) release. Mol Metab 3, 742–753 (2014).

35. Dokas J, et al. Conventional knockout of Tbc1d1 in mice impairs insulin- and AICAR-stimulated glucose uptake in skeletal muscle. Endocrinology 154, 3502–3514 (2013).

36. Szekeres F, et al. The Rab-GTPase-activating protein TBC1D1 regulates skeletal muscle glucose metabolism. Am J Physiol Endocrinol Metab 303, E524–533 (2012).

37. McDonagh B, Sakellariou GK, Jackson MJ. Application of redox proteomics to skeletal muscle aging and exercise. Biochem Soc Trans 42, 965–970 (2014).

38. Chrostek A, et al. Rac1 is crucial for hair follicle integrity but is not essential for maintenance of the epidermis. Molecular and cellular biology 26, 6957–6970 (2006).

39. Rao P, Monks DA. A tetracycline-inducible and skeletal muscle-specific Cre recombinase transgenic mouse. Dev Neurobiol 69, 401–406 (2009).

40. Pal R, Basu Thakur P, Li S, Minard C, Rodney GG. Real-time imaging of NADPH oxidase activity in living cells using a novel fluorescent protein reporter. PLoS One 8, e63989 (2013).

41. Knudsen JR, Henriquez-Olguin C, Li Z, Jensen TE. Electroporated GLUT4-7myc-GFP detects in vivo glucose transporter 4 translocation in skeletal muscle without discernible changes in GFP patterns. Exp Physiol, (2019).

