## Supplementary material for "Cytosolic ROS production by NADPH oxidase 2 regulates muscle glucose uptake during exercise": Supp data

Carlos Henríquez-Olguin *et al.*

**Supplementary Information**

**Figure S1. Similar abundance of antioxidant enzymes in *ncf1** muscles.** A) Total proteins levels of p38 MAPK, ERK 1/2, alpha2 AMPK, ACC, TBC1D1 and Coomassie staining as loading control in human muscle lysates. B) Antioxidants proteins from tibialis anterior muscle lysates from both *ncf1** and WT mice (n=13-14). C and D) cellular signaling for B) t-test was performed for statistical analysis, For C-D) Two-way ANOVA was performed to test for effects of exercise (Exer) genotype (Geno), and interaction (Int), followed by Tukey’s *post hoc* test to correct for multiple comparisons. Individual values and mean ± SEM are shown.

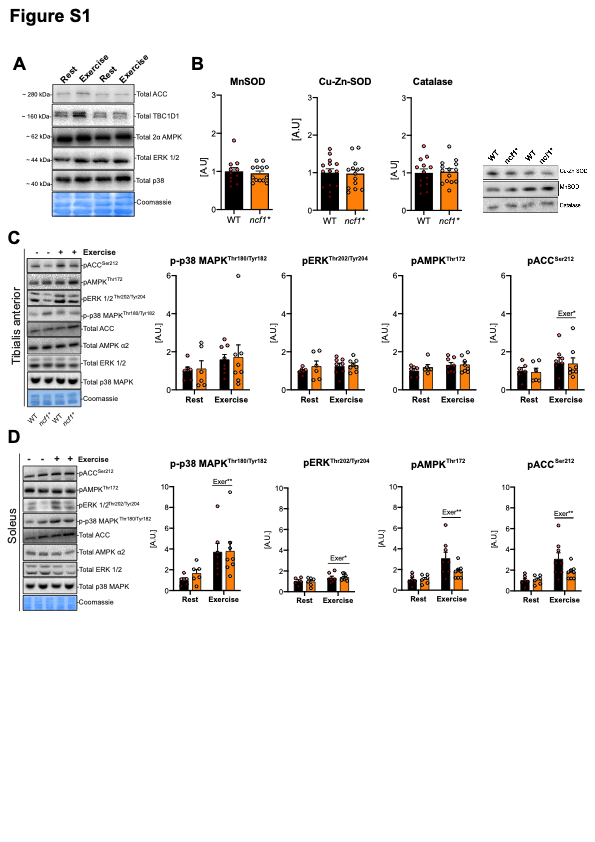

**Figure S2. Total protein levels of exercise-responsive kinases are similar in WT and *ncf1** muscles.** Protein abundance of p38 MAPK, ERK 1/2, alpha2 AMPK, and ACC in A) quadriceps (Quad) B) soleus (SOL) and tibialis anterior (TA) from WT and *ncf1** mice (n=14 per group). Individual values and mean ±SEM are shown.

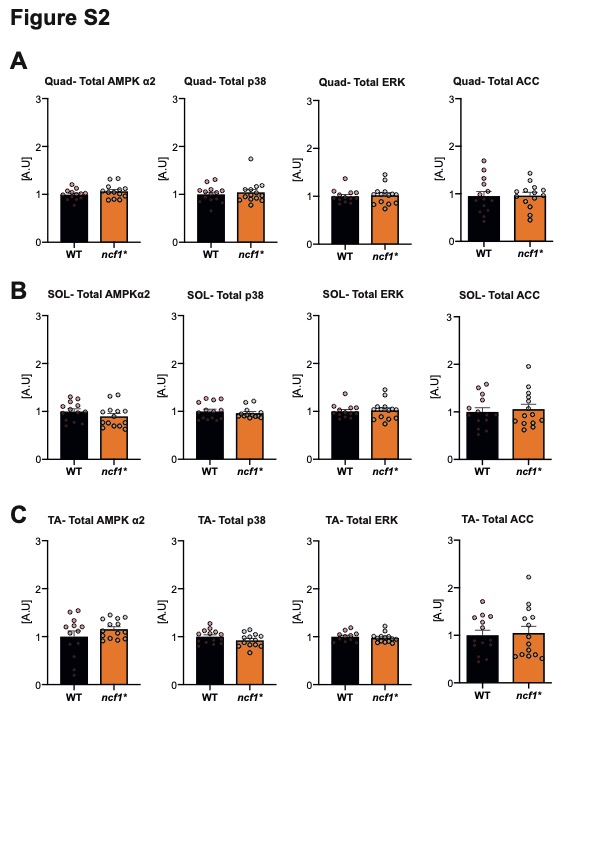

**Figure S3. Genetically-encoded biosensors are able to measure subcellular redox changes in muscle fibers.** A) Graphical description of the work flow for genetically encoded redox biosensors (the procedures are described in the methods section) B) Representative image and C) line-profile of the Mito-roGFP2-Orp1 and TMRE+ fluorescence in live flexor digitorum brevis (FDB) fibers. D) Mito-roGFP2-Orp1 oxidation by H_2_O_2_ (0.5 mM) stimulation in live FDB fibers. E) p47roGFP oxidation/reduction in response to H_2_O_2_ (0.5 mM) and DTT (0.5 mM). F) p47roGFP oxidation induced by electrical stimulation in FDB Rac1 imKO fibers (*n*=5). G) DCFH oxidation after 20 min of exercise in WT and Rac1 imKO Tibialis Anterior muscle (*n*=5-4). H) DCFH oxidation under resting conditions in WT and Rac1 imKO Tibialis Anterior muscle. I) Redox-regulated proteins in FDB fibers are similar between WT and Rac1 deficient muscles (*n*=4-5). One- way (for E, G, I) and Two-way ANOVA (for F) were performed to test for effects of exercise (Exer) genotype (Geno), and interaction (Int), Individual values and mean ± SEM are shown. # denotes P< 0.05 compared to the WT group.

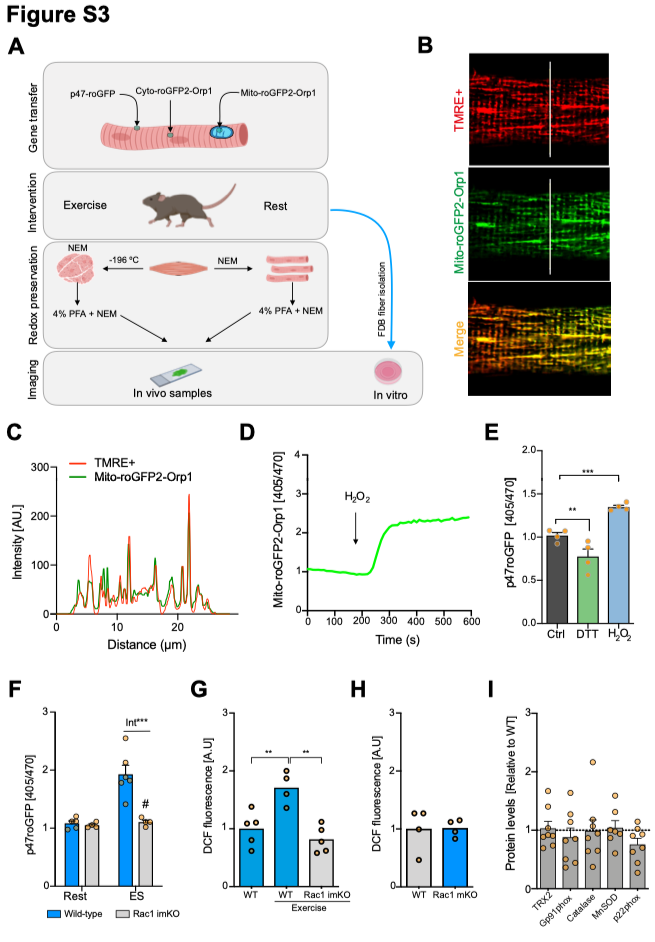

**Figure S4. Body weight and composition of *ncf1** mice.** A) Similar body weight despite tendency to B) lower body fat, C) higher lean mass and D) equal oxygen consumption in *ncf1** compared to WT mice. Unpaired t-test (F) and two-way ANOVA were performed to test for effects of exercise (Exer) genotype (Geno), and interaction (Int), followed by Tukey’s *post hoc* test to compare sub-groups. Individual values and mean ± SEM are shown.

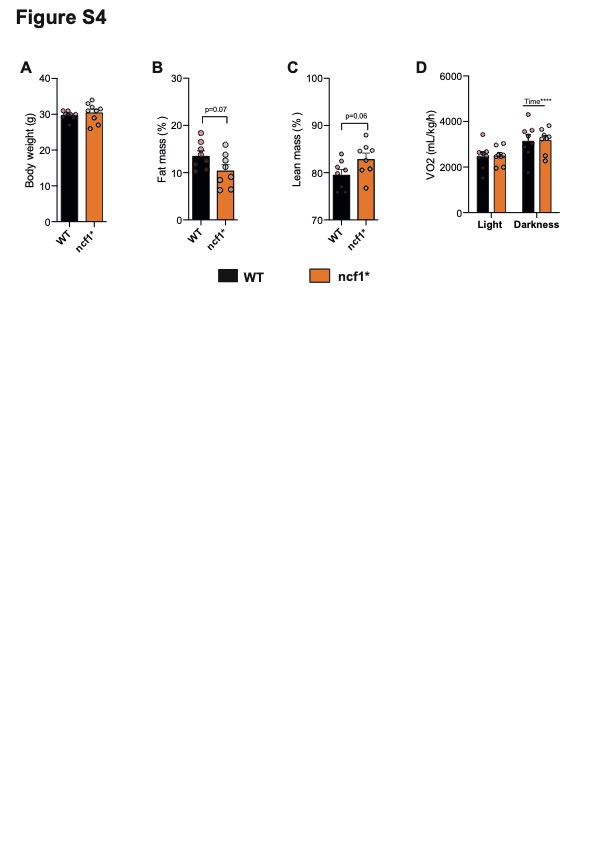

**Figure S5. Similar content of mitochondrial oxidative phosphorylation complex proteins in *ncf1** compared to WT muscles.** Quantification graphs of total protein showed figure 3E-F. A) quadriceps (n=12-13) and B) soleus muscle (n=13-14). Individual values and mean ± SEM are shown.

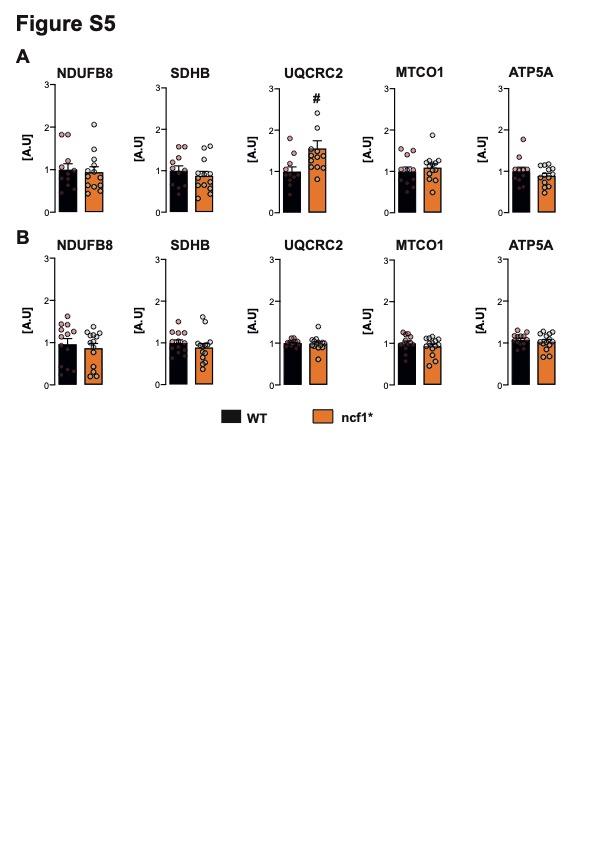

**Figure S6. Similar abundance of endogenous glucose handling-related proteins in *ncf1** and WT muscles.** Quantification graphs of total protein showed in figure 3B-D. A) quadriceps B) soleus, and C) tibialis anterior (TA) muscles. Individual values and mean ± SEM are shown

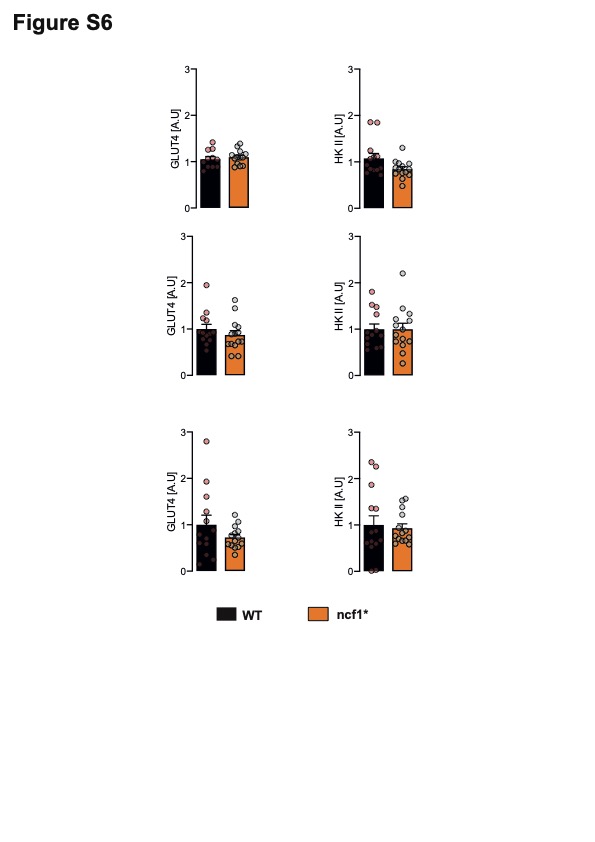

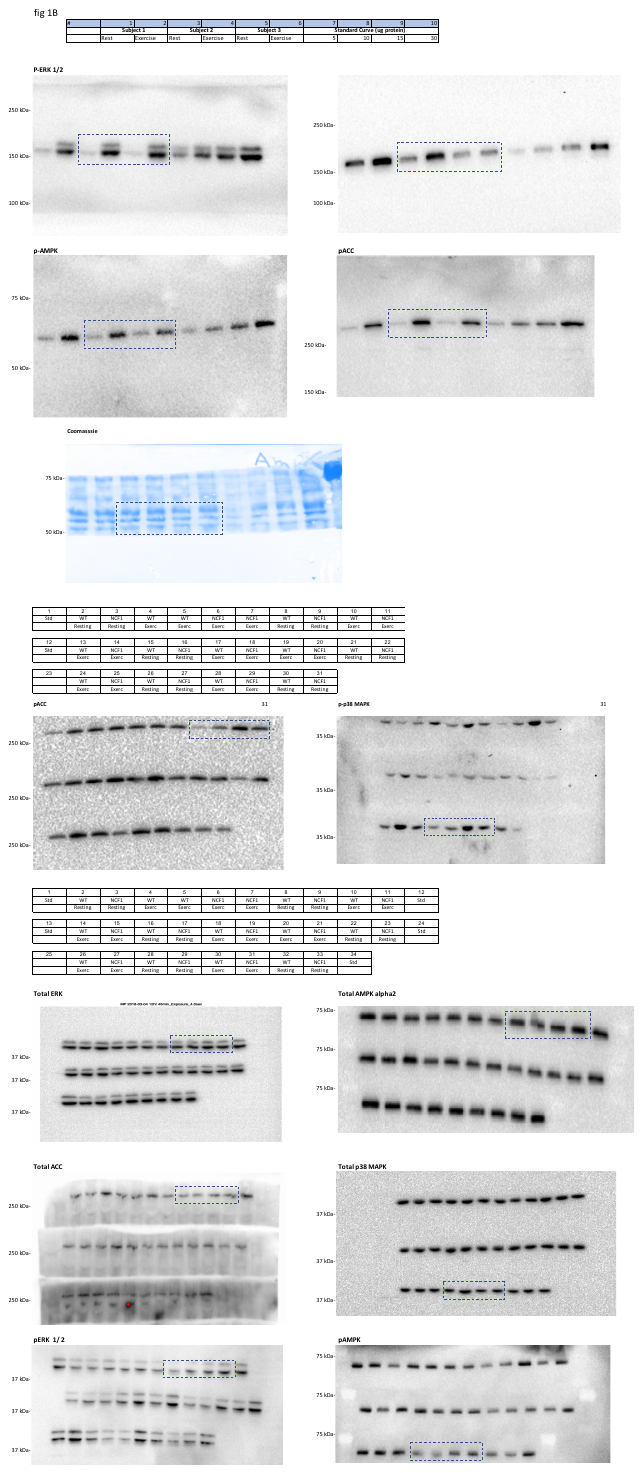
**Supplementary Figure 7: Full scans of western blots.**

**Supplementary Figure 7: Full scans of western blots (continuation)**

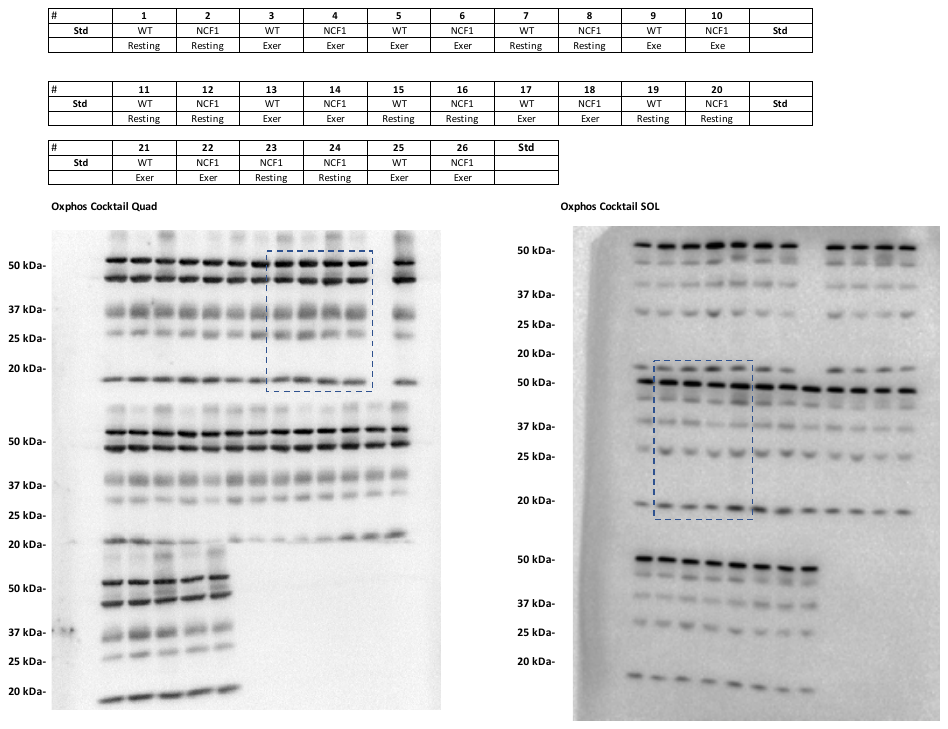

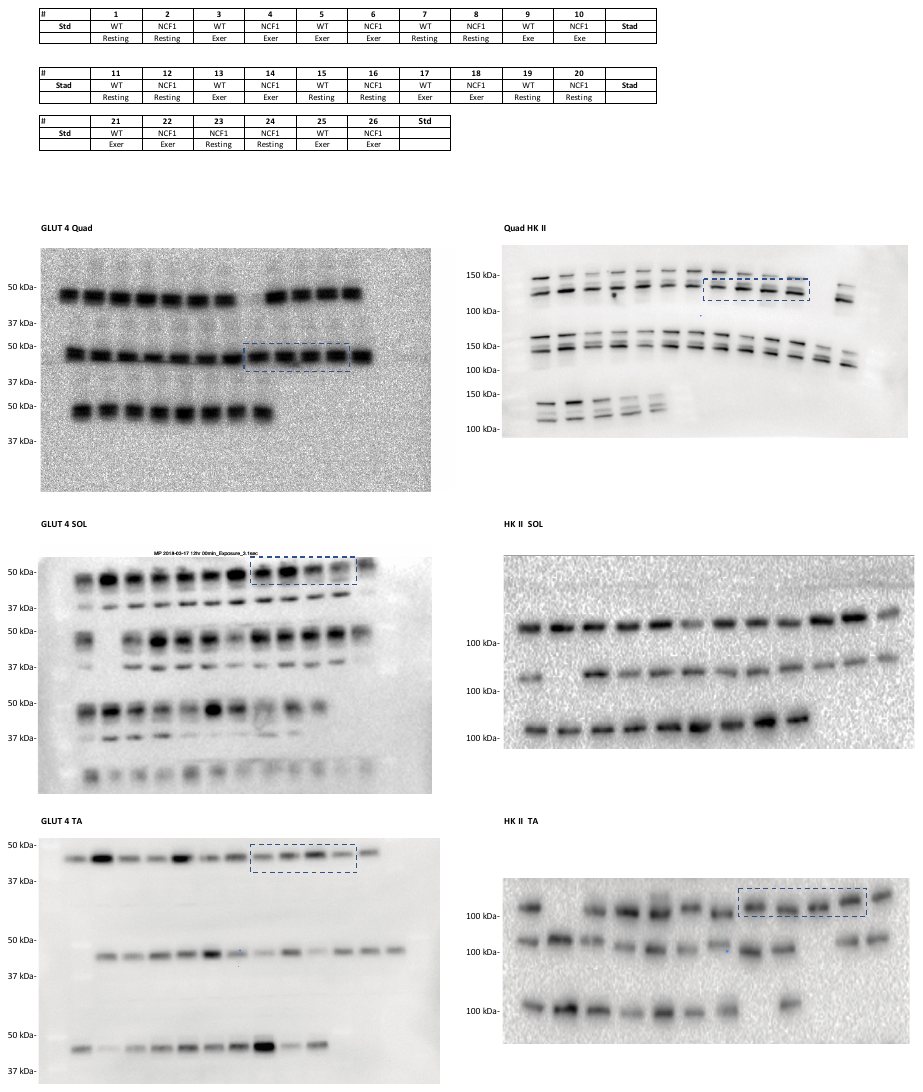
**Supplementary Figure 7: Full scans of western blots (continuation)**
